# Structural covariance networks are coupled to expression of genes enriched in supragranular layers of the human cortex

**DOI:** 10.1101/163758

**Authors:** Rafael Romero-Garcia, Kirstie J Whitaker, František Váša, Jakob Seidlitz, Maxwell Shinn, Peter Fonagy, Raymond J Dolan, Peter B Jones, Ian M Goodyer, the NSPN Consortium, Edward T Bullmore, Petra E Vértes

## Abstract

Complex network topology is characteristic of many biological systems, including anatomical and functional brain networks (connectomes). Here, we first constructed a structural covariance network (SCN) from MRI measures of cortical thickness on 296 healthy volunteers, aged 14-24 years. Next, we designed a new algorithm for matching sample locations from the Allen Brain Atlas to the nodes of the SCN. Subsequently we use this to define, transcriptomic brain networks (TBN) by estimating gene co-expression between pairs of cortical regions. Finally, we explore the hypothesis that TBN and the SCN are coupled.

TBN and SCN were correlated across connection weights and showed qualitatively similar complex topological properties. There were differences between networks in degree and distance distributions. However, cortical areas connected to each other within modules of the SCN network had significantly higher levels of whole genome co-expression than expected by chance.

Nodes connected in the SCN had significantly higher levels of expression and co-expression of a Human Supragranular Enriched (HSE) gene set that are known to be important for large-scale cortico-cortical connectivity. This coupling of brain transcriptome and connectome topologies was largely but not completely related to the common constraint of physical distance on both networks.

## INTRODUCTION

Many natural systems have a complex network topology. Nervous systems form anatomical networks with graph-theoretically non-random properties like small-worldness, high-degree hubs and modules (Bullmore and Sporns 2009). Brain networks conserve these topological features from the micro (synaptic) scale of *C. elegans* to the macro scale of whole human brain connectomes derived from statistical analysis of human magnetic resonance imaging (MRI) data (Fornito et al. 2016). A similar topological profile has also been described for networks of gene expression, in which each node is a gene and each edge represents the strength of correlation or co-expression of genes between samples collected in different tissue types or across different individuals or different time-points (Belcastro et al. 2011).

Here we define a novel kind of brain network whose nodes represent brain regions and whose links are based on pairwise correlations in whole-genome gene expression across brain regions. We then investigate the relationship between this transcriptomic brain network (TBN) and a structural covariance network (SCN) constructed from MRI data on 296 healthy participants, representing the human brain anatomical connectome.

The method of structural covariance describes the existence of positively correlated brain regional anatomical measurements – such as cortical thickness or volume – between pairs of brain regions (Wright et al. 1999). The structural covariance matrix of a group of subjects is defined by estimating the inter-regional correlation of cortical thickness between all possible pairs of regions defined by an anatomical parcellation. The covariance matrix can be thresholded to construct a binary or weighted graph of anatomical covariation, which putatively represents anatomical connectivity (He et al. 2007). Structural covariance networks are replicable, heritable, and representative of disease-related changes in topology (Alexander-Bloch et al. 2013). However, the neurobiological substrate of inter-regional structural covariation remains poorly understood. There is some evidence that structural correlation is related to anatomical connectivity between regions; but MRI networks based on inter-regional correlation of cortical thickness (CT) are only moderately similar to DTI networks based on tractographic analysis of white matter projections between cortical areas (Gong et al. 2012).

Gene expression in all regions of human cortex can be estimated using the transcriptomic dataset made publicly available by the Allen Institute for Brain Science (AIBS). The AIBS used *in situ* hybridization to produce the first comprehensive genome-wide atlas of mRNA expression that can be mapped into MNI anatomical space (Hawrylycz et al. 2012). Previous analysis of these data has demonstrated that the transcriptional profile of cortical tissue is fairly homogenous compared to variations between cortex and subcortex (Hawrylycz et al. 2012); and the relatively subtle variations of cortical gene expression are associated with cytoarchitectonic gradation across the cortex (Chen et al. 2013). Cortical gene expression has also been shown to correlate with resting state functional connectivity (Hawrylycz et al. 2015) and morphometric similarity (Seidlitz et al., 2017). Functional connectivity and morphometric similarity were specifically related to expression of a set of human supragranular expressed (HSE) genes that are highly expressed in the supragranular layers of human cortex (lamina 2, 3), where most cortico-cortical projections originate (Krienen et al. 2016; Vértes et al. 2016). Other studies have shown that functional connectivity between regions is associated with co-expression of genes enriched for ion channel and synaptic ontology terms (Hawrylycz et al. 2015; Richiardi et al. 2015).

On this basis, we hypothesized that putative metrics of anatomical connectivity of human cortex – such as inter-regional correlations of cortical thickness – should be related to gene co-expression, and specifically co-expression of HSE genes. In what follows, we first develop a new method for matching samples from the AIBS to MRI regions of interest in the native space of each postmortem AIBS brain. On this basis we construct a transcriptomic brain network (TBN) and describe its graph theoretical properties, as well as comparable properties of the structural covariance network (SCN; connectome). Finally, we explore the relationship between the two networks, testing the specific hypotheses: (i) that regions connected as part of the same modules of the SCN had higher levels of whole genome and HSE gene co-expression than regions in different modules; (ii) that regions connected in the SCN had higher levels of HSE gene co-expression than regional nodes that were not anatomically connected; and (iii) that the relationships between structural correlation and gene co-expression were not entirely attributable to common constraints of physical connection distance on both these spatially embedded networks.

## METHODS

### Participants

2135 healthy young people in the age range 14-25 years were recruited from schools, colleges, NHS primary care services and direct advertisement in north London and Cambridgeshire. This primary cohort was stratified into 5 contiguous age-related strata: 14-15 years inclusive, 16-17 years, 18-19 years, 20-21 years, and 22-25 years. Recruitment within each stratum was evenly balanced for sex and ethnicity and satisfied exclusion criteria including any history of treatment for psychiatric disorder or drug or alcohol dependence, or any history of neurological disorder, head injury or learning disability. A demographically balanced cohort of N=300 was sub-sampled from the primary cohort for structural MRI scanning in one of the following sites: (1) Wellcome Trust Centre for Neuroimaging, London; (2) Medical Research Council Cognition and Brain Sciences Unit and (3) Wolfson Brain Imaging Centre, Cambridge. Four MRI scans were excluded due to insufficient quality, resulting in a final MRI sample of 296 subjects (19.11 ± 2.93 years [mean ± standard deviation], 148 females).

Written informed consent was provided by all participants as well as written parental assent for participants less than 16 years old. The study was ethically approved by the National Research Ethics Service and was conducted in accordance with NHS research governance standards.

### MRI data acquisition

Structural MRI scans were acquired on one of three identical 3T MRI systems (Magnetom TIM Trio, Siemens Healthcare, Erlangen, Germany; VB17 software version) equipped with a standard 32-channel radio-frequency (RF) receive head coil and RF body coil for transmission. The multi-parametric mapping (MPM) protocol (Weiskopf et al. 2013) yields 3 multi-echo fast low angle shot (FLASH) scans with variable excitation flip angles. By appropriate choice of repetition time (TR) and flip angle α, acquisitions were predominantly weighted by T1 (TR=18.7ms, α=20°), Proton Density (PD) or Magnetization Transfer (MT) (TR=23.7ms, α=6°). Multiple gradient echoes were acquired with alternating readout polarity at six equidistant echo times (TE) between 2.2 and 14.7 ms for the T1-weighted and MT-weighted acquisitions and at 8 equidistant TE between 2.2 ms and 19.7 ms for the PD-weighted acquisition. The resulting three mean images were used to calculate the parameter maps of the apparent longitudinal relaxation rate R1, the MT saturation, and the effective proton density PD* using previously developed models describing the image intensity of FLASH scans (Helms, Dathe, and Dechent 2008; Helms, Dathe, Kallenberg, et al. 2008; Weiskopf et al. 2011). Here, only R1 quantitative maps were used.

Quantitative R1 maps were determined from the apparent R1 maps by correcting for local RF transmit field inhomogeneities and imperfect RF spoiling (Preibisch and Deichmann 2009). This approach was adapted to the FLASH acquisition parameters used here. RF transmit field maps were calculated from the 3D EPI acquisition and corrected for off-resonance effects (Lutti et al. 2012).

Other acquisition parameters were: 1 mm^3^ voxel resolution, 176 sagittal slices, field of view (FOV) = 256 x 240 mm, parallel imaging using GRAPPA factor 2 in phase-encoding (PE) direction (AP), 6/8 partial Fourier in partition direction, non-selective RF excitation, readout bandwidth BW = 425 Hz/pixel, RF spoiling phase increment = 50°. A pilot study demonstrated satisfactory levels of between-site reliability in MPM data acquisition (Weiskopf et al. 2013).

### MRI reconstruction, cortical parcellation and SCN construction

We used a standard automated processing pipeline (FreeSurfer v5.3) for skull stripping, tissue classification, surface extraction and cortical parcellation (http://surfer.nmr.mgn.harvard.edu) applied to longitudinal relaxation rate (R1) maps (R1=1/T1). Cortical thickness (CT) measurements were estimated by reconstructing the pial surface and the boundary between grey matter and white matter (Dale and Sereno 1993; Dale et al. 1999) and measuring the distance between these surfaces; see **Figure 1A**. Errors in grey/white matter boundary reconstruction were corrected by manual editing to improve cortical thickness estimation. Four scans did not pass quality control for accurate cortical segmentation and were excluded due to movement artefacts.

**Figure 1:**
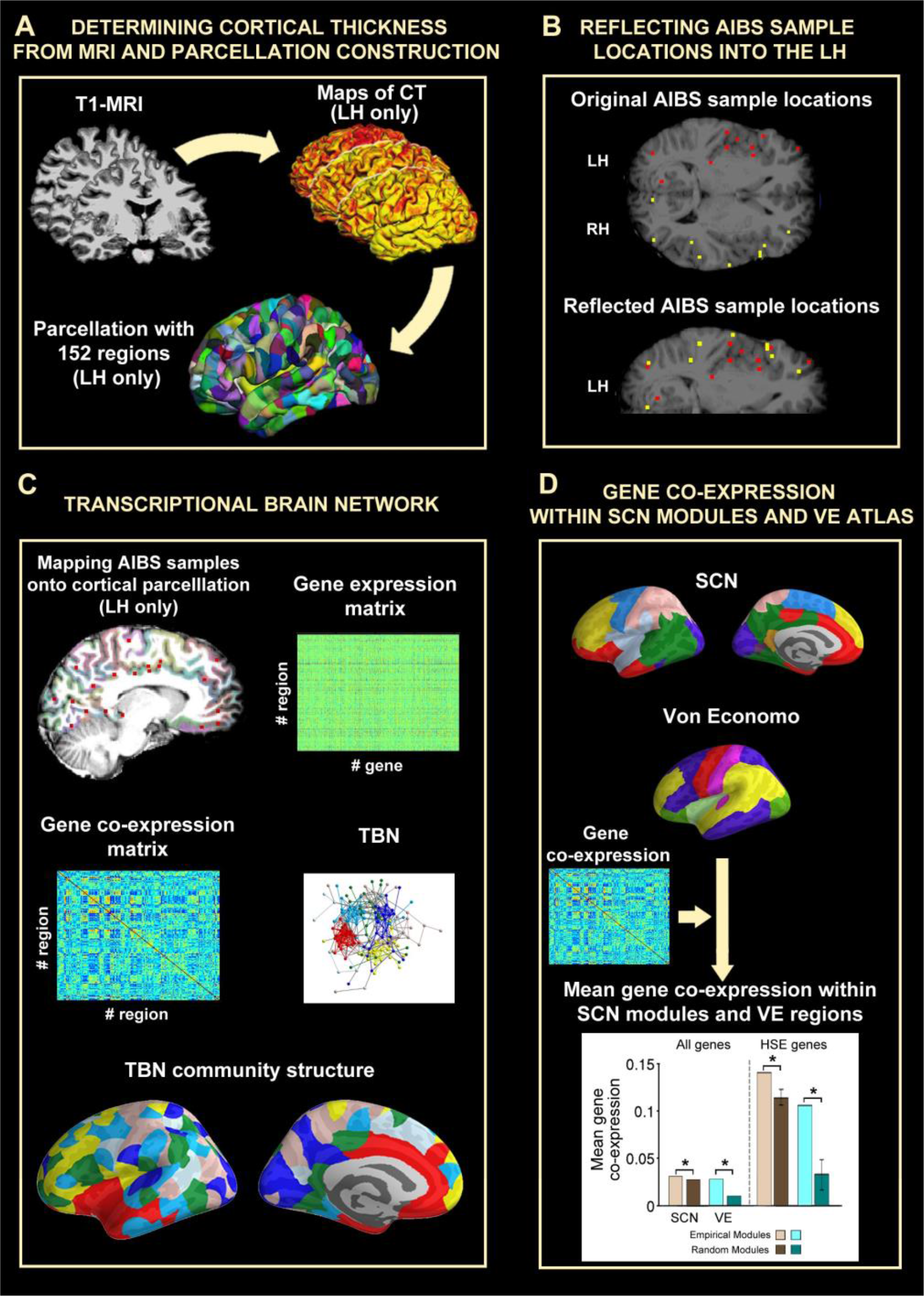
A schematic overview of data analysis. **A:** Cortical thickness was estimated from T1-weighted MRI scans on 296 healthy young participants. Left hemisphere (LH) was subdivided into regional nodes by a fine-grained parcellation of 152 nodes based on subdivision of Desikan-Killiany nodes. **B:** Reflection of samples located in the right hemisphere (RH, yellow dots) into the LH (red dots) to increase sample density. **C:** Regional gene expression from the AIBS Human Brain Atlas were mapped to the same cortical parcellation templates to estimate regional expression profiles and a co-expression matrix. The gene expression matrix was thresholded to construct a binary graph which had a modular community structure. **D:** Modules of the structural covariance network, and cytoarchitectonic classes defined by von Economo, were used to test the hypothesis that whole genome co-expression and HSE gene co-expression were greater between nodes in the same topological module or cytoarchitectonic class.

Due to under-sampling of the right cerebral hemisphere in the gene expression data (see below; Hawrylycz et al., 2012) we focused attention on the left hemisphere of both the MRI and gene expression data. We used a high-resolution parcellation of the left hemisphere that comprised 152 cortical regions, each with an approximate surface area of 500 mm^2^. Consequently, MRI data corresponding to the right hemisphere was discarded. This cortical scheme was created by sub-dividing the anatomically defined regions in the Desikan-Killiany atlas (DK, Desikan et al., 2006) for the FreeSurfer surface template (fsaverage) (Romero-Garcia et al. 2012); see **Figure 1A**. R1-weighted images of each subject were co-registered to the FreeSurfer surface template using rigid transformations to obtain the transformation matrix *T*_*R*1_. Inverse transformations 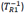were used to warp the parcellation scheme from standard stereotactic space to the space of each individual R1-weighted image. CT values were averaged across all vertices included in each cortical parcel. This process was repeated for each subject resulting in a (296 × 152) matrix of cortical thickness estimates at each of 152 regions for each of 296 participants.

SCN construction relies on the identification of spatial patterns of morphometric similarities between brain regions within a group of subjects. The inter-regional correlations in cortical thickness were estimated to construct a (152 × 152) structural correlation matrix. A hard threshold was applied to this matrix so that an arbitrary percentage (connection density) of the most strongly positive CT correlations were retained as non-zero elements in a binary adjacency matrix or, equivalently, edges between regional nodes in a graph of the structural covariance network. Brain networks were visualized using BrainNet viewer (Xia et al. 2013) (http://www.nitrc.org/projects/bnv/).

### AIBS gene expression dataset

We used the whole genome expression atlas of the adult human brain created by the Allen Institute for Brain Science (http://human.brain-map.org) (Hawrylycz et al. 2012, 2015). The AIBS dataset includes samples from post-mortem brains of six donors (3 Caucasian, 2 African-American, 1 Hispanic) aged 24-57 years. Microarray analysis was used to measure expression of all genes in the genome at each of multiple cortical and subcortical locations in each donor’s brain. As different cRNA hybridization probes were used to identify the expression level of the same gene, expression values from multiple probes were averaged for each gene. Probes that were not matched to gene symbols in the AIBS data were excluded, resulting in 20,737 gene expression values that were evaluated in 3702 brain samples. Due to the similarity of gene expression between hemispheres (Hawrylycz et al. 2012; Pletikos et al. 2014), the AIBS only sampled two of the six donors in the right hemisphere. Therefore, we only focused on the left hemisphere in further analyses. However, given the strong expression similarities between hemispheres and in order to increase the number of gene expression samples per cortical region, all right hemisphere samples were reflected to their contralateral position into the left hemisphere (**Figure 1B**). Consequently, final gene expression profile of each region was considered including both its ipsilateral samples (from all six donors) and its contralateral samples (from the two donors whose right hemisphere was sampled).

### Matching MRI parcellation with samples location

We develop a new method to map the anatomical structure associated to each tissue sample by using the MRI data provided by the AIBS for each donor. T1-images of the six donors were processed following the FreeSurfer pipeline. The high-resolution parcellation with 152 cortical regions in the left hemisphere, used in the analysis of the MRI data, was warped from fsaverage space to the surface reconstruction of each AIBS donor’s brain. The surface-based parcellation of each donor’s brain was transformed into a volumetric parcellation that covered the whole cortex and extended 2mm into the subjacent white matter to include those cortical samples that had been excluded due to registration misalignment; **Figure 1C**. 91% of the total AIBS gene expression cortical samples were located within the resulting volumetric parcels; **Figure S1** shows the number of AIBS samples covered by each cortical region. Gene expression values for each cortical parcel were estimated as the median normalized gene expression over all six donors and compiled to form a (152 × 20,737) matrix representing the expression of each of 20,737 genes at each of 152 left hemisphere cortical regions; **Figure 1C**. Code used to estimate the gene expression levels in each matrix is available at: https://github.com/RafaelRomeroGarcia/geneExpression_Repository.

### Transcriptional brain network construction

The (152 x 152) gene co-expression matrix was estimated by the pairwise Pearson’s correlations between whole genome expression profiles in each possible pair of cortical regions (**Figure 1C**). A hard threshold was applied to this matrix so that an arbitrary percentage (connection density) of the most strongly positive transcriptional correlations were retained as non-zero elements in a binary adjacency matrix or, equivalently, edges between regional nodes in a graph of the transcriptional brain network (TBN).

### Human supragranular enriched (HSE) genes

The human supragranular enriched (HSE) gene list comprises 19 genes: *BEND5, C1QL2, CACNA1E, COL24A1, COL6A1, CRYM, KCNC3, KCNH4, LGALS1, MFGE8, NEFH, PRSS12, SCN3B, SCN4B, SNCG, SV2C, SYT2, TPBG* and *VAMP1* (Zeng et al. 2012). The normalized median expression profile of each HSE gene in each of 152 cortical regions was extracted from the whole genome transcription matrix.

### Network analysis and community structure

Global topological properties of the structural correlation and transcriptional networks were analyzed using the following graph theoretical measures: clustering coefficient (*C*_*p*_), path length (*L*_*p*_), local efficiency (*E*_*loc*_), global efficiency (*E*_*glob*_), small-worldness (σ) and assortativity (*a*). Such topological properties are strongly influenced by more fundamental features of the network, including the number of nodes, number of connections, and degree distribution. To control these effects, network measures for the empirical networks were compared with those for 100 randomized networks generated using a random rewiring process (Maslov and Sneppen 2002). At a nodal level, we estimated degree centrality as the number of edges connecting each node to the rest of the network. Global topological metrics were computed using the Brain Connectivity Toolbox (https://sites.google.com/site/bctnet/) (Rubinov and Sporns 2010).

High degree nodes, also known as hubs, are often densely inter-connected to form a rich club. The rich club coefficients (ϕ(*r*)) of a thresholded network were computed as the sum of edges within the subgraph defined by retaining only nodes with degree greater than an arbitrary threshold. The rich club metric was normalized (*r*_*norm*_(*k*)) by computing the ratio of the rich club coefficient of the real network (ϕ(*k*)) to the mean rich club coefficient of 100 randomized networks (ϕ_*rand*_(*k*)).

Modularity analysis was used to decompose the community structure of each network into a set of modules that each comprised a community of nodes that were densely connected to each other but sparsely connected to nodes in other modules (**Figure 1C**). Community partitioning was performed by maximizing the metric defined by Newman (2006) comparing the density of intra-modular connections to the density expected to occur by chance in a random network:

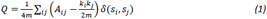

where *m* is the total number of edges in a network, *A*_*ij*_ is the adjacency matrix, *k*_*i*_ and *k*_*j*_ are the degree of nodes *i* and *j*, δ(*s*_*i*_, *s*_*j*_) is the Kronecker delta, and *s*_*i*_ and *s*_*j*_ are the communities of nodes *i* and *j*, respectively. We used a non-deterministic modularity algorithm which computes a local optimum by greedy optimization (Blondel et al. 2008). In order to detect a stable consensus community structure, this modularity decomposition algorithm was applied 1000 times. A (152 x 152) consensus matrix (M) was created defining each element *m*_*ij*_ as the number of times that node *i* and node *j* had been classified in the same module. Finally, the consensus matrix was used as an input for the community algorithm. The resulting output represented a stable modular structure of the original network (Kwak et al. 2009).

### Gene co-expression within modules derived from SCN and Von Economo

To assess the spatial overlap between SCN community partitions and gene co-expression, we averaged the gene co-expression values between each pair of regions located within the same SCN module. As this intra-modular gene co-expression is affected by trivial characteristics of the community partition, raw values were compared with those derived from random modular partitions. To test against appropriately designed surrogate data, 1000 pseudo-random communities were created by iteratively permuting the module labels associated to pairs of nodes located at the same distance from each module’s centroid. This procedure randomly shifts the position of the modules along the cortex without splitting apart the components of the module (**Figure S2**) (Bethlehem et al. 2017). The resulting null distribution of community partitions preserves the number and size of modules, as well as the spatial contiguity of the empirical community partition. The 95th percentile of the null distribution was used as a statistical threshold to retain or reject the null hypothesis of no significant gene co-expression within modules. **Figure 1D** illustrates the estimation of the mean gene co-expression within the two modular partitions considered in this paper: modules derived from the SCN, and the von Economo atlas. This atlas groups brain regions according to cytoarchitectonic criteria (von Economo 1929). Thus, the complete cortex was divided into: (i) granular cortex; primary motor/precentral gyrus, (ii) association cortex, (iii) association cortex, (iv) secondary sensory cortex, (v) primary sensory cortex. Due to their unique cytoarchitectonical features (vi) limbic regions (including entorhinal, retrosplenial, presubicular and cingulate) and (vii) insular cortex (which contains granular, agranular and dysgranular regions) were considered as separate modules (Vértes et al. 2016; Seidlitz et al. 2017; Vasa et al. 2017).

## RESULTS

### Topological and spatial properties of the transcriptomic brain network (TBN) and the structural covariance network (SCN)

Following thresholding to 10% edge density, nodes in both the TBN and SCN were part of a single connected component. Topologically, both networks were small-world (as defined by greater than random clustering combined with near random path length or global efficiency) with a modular community structure and a rich club of highly interconnected hub nodes (**Figure 2**).

**Figure 2:**
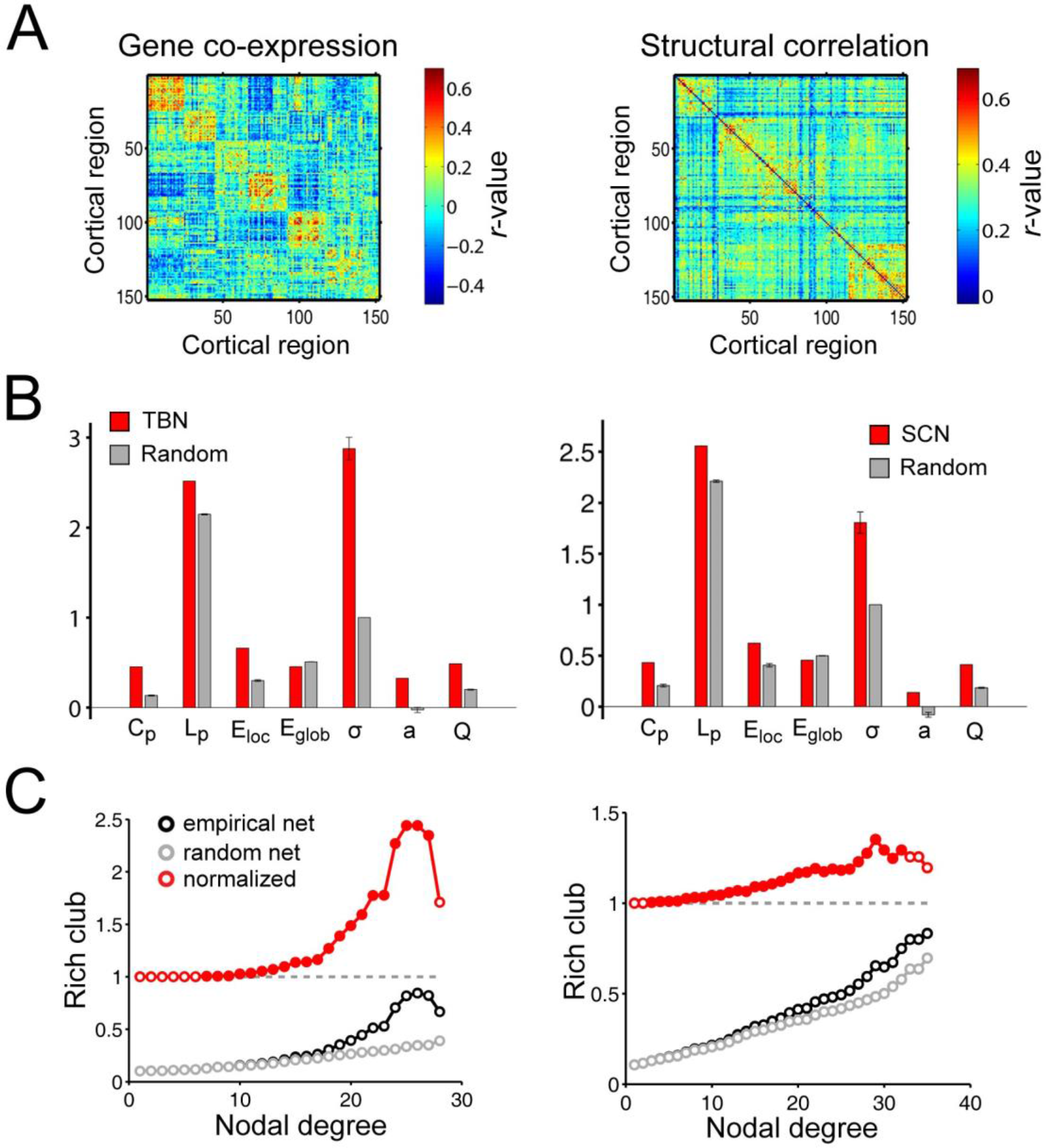
Global topology of gene co-expression (left) and structural covariance networks (right). **A**: Gene co-expression matrix and structural covariance matrix identically ordered in alignment with the modular community structure of the transcriptional network. B) Global topological metrics estimated in TBN, SCN and comparable random networks: Cp = clustering coefficient; Lp = path length; Eloc = local efficiency; Eglob = global efficiency; σ = small-world; a = assortativity; Q = modularity. Error bars represents the standard deviations. **C**: Rich club coefficient curves for TBN, SCN and random networks.

The degree distribution of both networks was fat-tailed compared to a random graph (**Figure 3A**, left panel), the SCN having more highly-connected hubs than the TBN. In both networks, the distance distribution was skewed towards shorter distance, compared to a random graph, but there was a tail of long distance connections that was particularly prominent in the TBN (**Figure 3A**, right panel). Structural covariation strength and gene co-expression strength both decreased monotonically as a function of increasing physical distance between nodes (R^2^=0.20 and R^2^=0.15, respectively; **Figure 3B**, first and second panel). Akaike’s information criterion (AIC) demonstrated that an exponential function of distance provided a better fit than a linear function for both the structural covariance (*AIC*_*linear*_ = −0.33 and *AIC*_*exponential*_ = −0.55) and gene co-expression (*AIC*_*linear*_ = 5.13 and *AIC*_*exponential*_ = 5.11). Gene co-expression and structural covariance were significantly correlated (R^2^=0.05, P<10^-10^; **Figure 3B**, third panel). This relationship remains significant after correcting for inter-regional distance (R^2^=0.01, P<10^-10^; **Figure S3**). Nodal degree was significantly related to the average Euclidean distance between connected nodes in the SCN (R^2^=0.51, P<10^-10^; **Figure 3C**, first panel), i.e., high degree hubs had more long-distance connections to other nodes; but there was no relationship between distance and degree in the TBN (R^2^<10^-3^, P=0.86; **Figure 3C**, second panel). Nodal degree was not correlated between the structural covariance and transcriptional networks (R^2^=0.01, P=0.19; **Figure 3C**, third panel), i.e., the high degree hubs of the SCN did not correspond to the hubs of the TBN.

**Figure 3:**
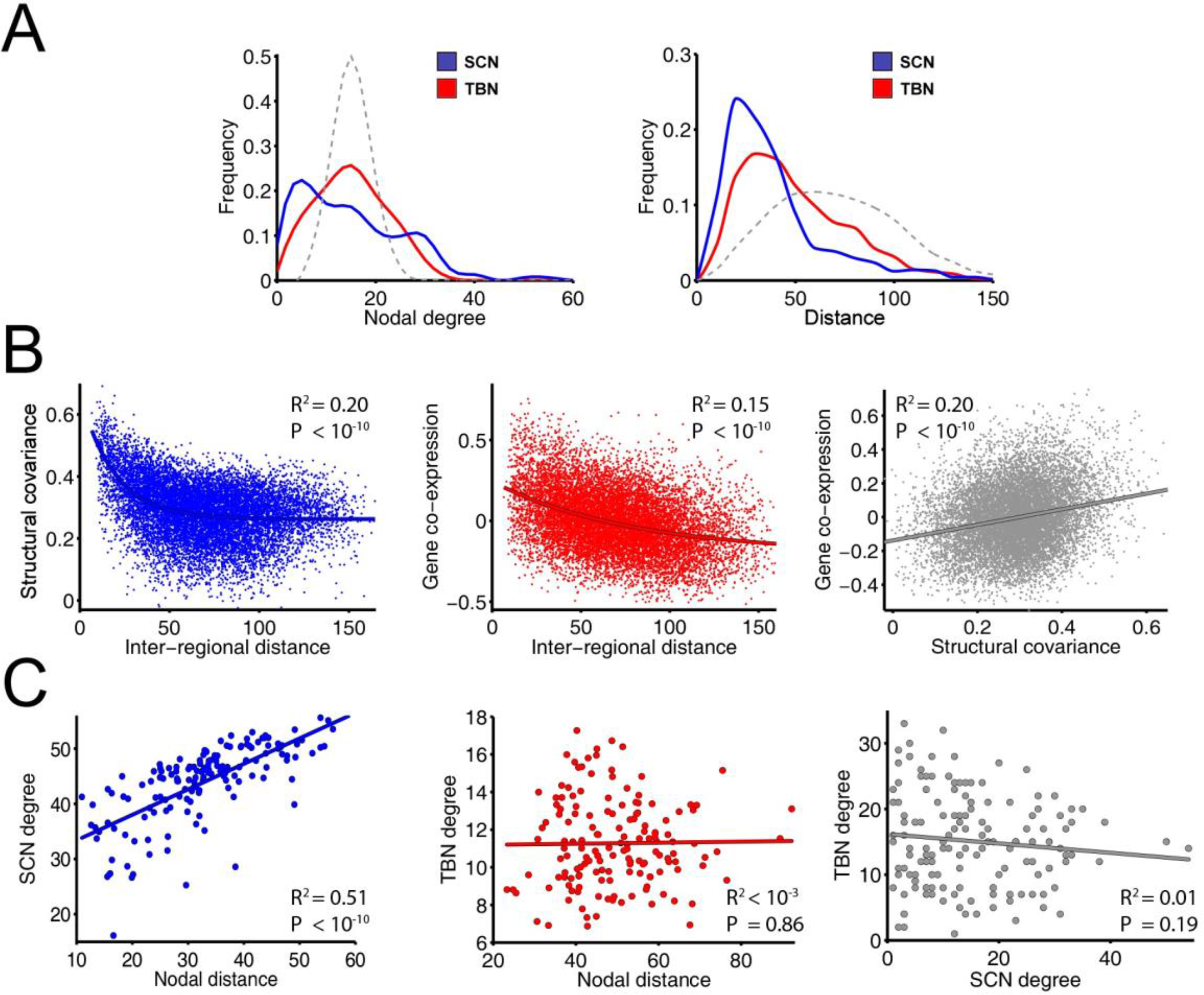
Nodal topology and connection distance of structural covariance network (SCN) and transcriptional brain network (TBN). **A**. Degree distribution of both networks (solid lines) and a comparable random network (dashed line). Connection distance distribution of both networks (solid lines) and a comparable random network (dashed line). **B.** Effect of inter-regional distance on structural covariance (left); effect of inter-regional distance on gene co-expression (middle); and gene co-expression versus structural covariance (right). **C.** Degree versus connection distance in the SCN (left); degree versus connection distance in the TBN (middle); and nodal degree in TBN versus nodal degree in SCN.

### Whole genome co-expression in relation to the modular community structure of the SCN

The spatial location of TBN modules was partially overlapping with the community structure derived from the structural covariance network (**Figure 4A**). Gene co-expression was significantly higher between regional nodes that belonged to the same module of the SCN than between nodes that belonged to randomly defined modules that preserved the number, size and spatial contiguity of the SCN modules (**Figure 4B**, left panel; P<0.005). This result was replicated for different SCN costs (5%, 10% and 15%) and modularity resolution parameters (γ=1 and γ=2) (all P<0.05; **Figure S4**). Gene co-expression was also significantly increased within regions grouped according to the cytoarchitectonic criteria of von Economo (P<0.02; **Figure 4B**, right panel; (von Economo 1929)). In other words, gene co-expression was increased between regions that belonged to the same cytoarchietctonic class.

**Figure 4:**
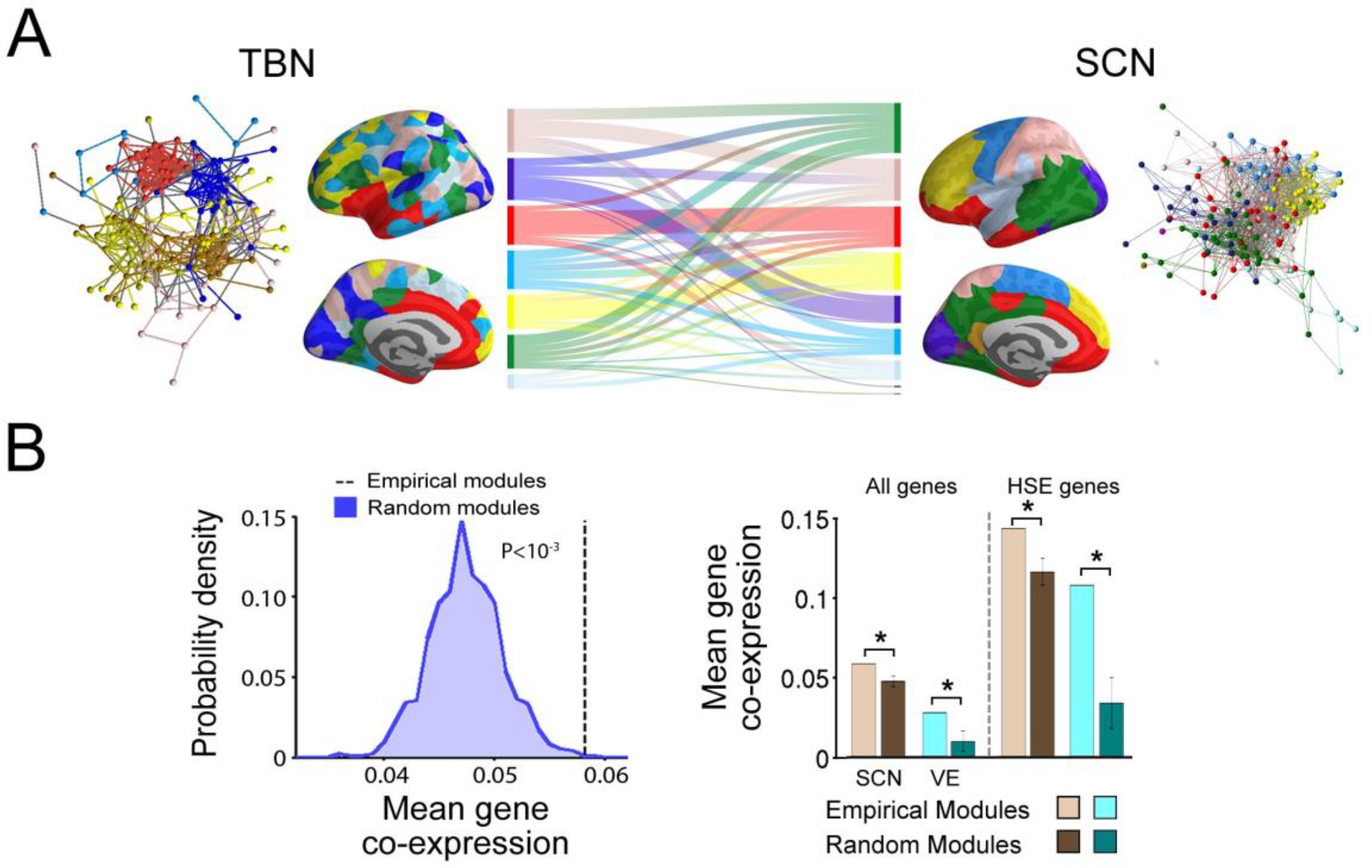
Community structure of the structural covariance network and transcriptomic brain network. **A:** Modular decomposition of the whole genome transcriptional brain network (7 modules; left); alluvial diagram showing how regions were aligned to the same or different modules in the two networks (middle); and modular decomposition of the structural covariance network (9 modules; right). **B**: Distribution of whole genome co-expression between regions assigned to the same random modules compared to the whole genome co-expression between regions assigned to the same modules of the empirical SCN (left). Co-expression of the whole genome and the HSE genes within each module of the SCN, within each cytoarchitectonic class of the von Economo (VE) atlas, and within comparable random modules. Error bars represents the standard deviations.

### HSE gene expression and structural covariation

Expression of HSE gene set was explored due to its particular transcriptional profile in supragranular layers of the human cortex and its association with long-range cortico-cortical connectivity (Zeng et al. 2012). HSE genes showed a heterogeneous expression pattern across the cortex that partially overlaps with the anatomical patterning of nodal degree in the structural covariance network (**Figure 5A**). HSE genes were over-expressed (compared with the whole brain mean expression level) in von Economo classes (i) granular, primary motor, (ii) association cortex and (iii) association cortex; whereas HSE were under-expressed in classes (iv) secondary sensory cortex, (v) primary sensory cortex, (vi) limbic regions and (vii) insular cortex (**Figure 5B;** P<0.05, FDR-corrected). Nodal degree in the SCN was significantly correlated with HSE gene expression (R^*2*^=0.14; P<10^−5^; **Figure 5C**). SCN nodal degree was higher in von Economo classes 1,2,3 and 4 than in classes 5,6 and 7 (**Figure 5D**).

**Figure 5:**
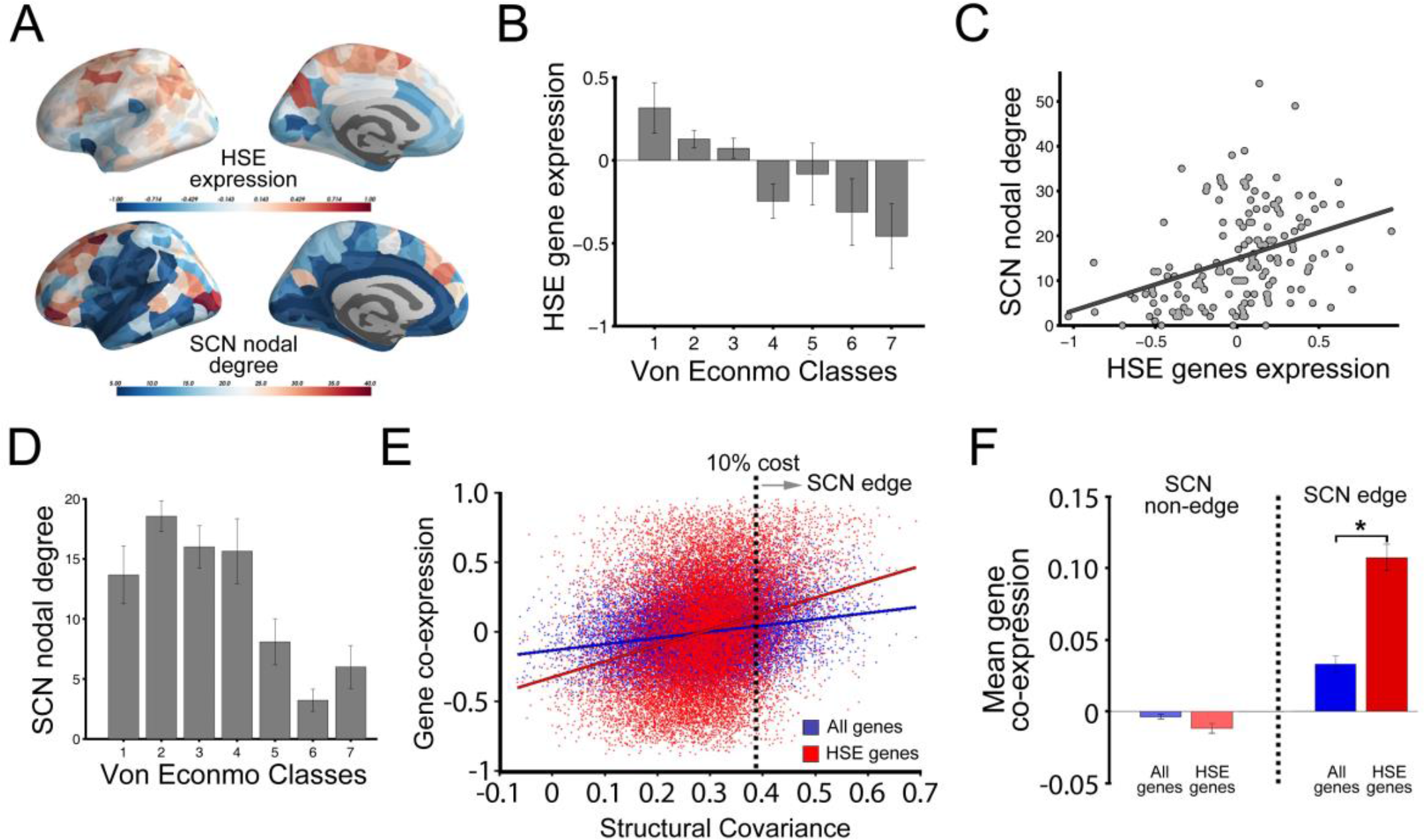
Structural covariance and HSE gene co-expression. **A:** HSE gene expression (top) and SCN nodal degree (bottom). **B:** HSE gene expression in different cytoarchitectonic classes defined by von Economo atlas. **C:** Nodal degree in structural covariance network versus HSE gene expression. **D:** Mean nodal degree in different cytoarchitectonic classes defined by von Economo atlas. **E:** Co-expression (whole genome, blue; or HSE genes only, red) versus structural covariance. Dashed vertical line indicates the threshold value of structural covariance used to define a binary graph of edges and non-edges. **F:** Co-expression (whole genome, blue; or HSE genes only, red) for SCN non-edges (unconnected regions) and SCN edges.

Correlation between structural covariance and gene co-expression was higher for HSE genes (R^*2*^=0.09, P<10^−10^) than when the complete genome was considered (R^*2*^=0.06, P<10^−10^; permutation test) and this difference was statistically significant (*F*_1,11475_=130.5, P<10^−4^, after regressing out the effect of the distance; **Figure 5E**). Gene co-expression between connected regions in the structural covariance network (SCN edges) was higher for HSE genes (mean Pearson *r* = 0.11) than for the complete genome (mean Pearson *r* = 0.03) and this difference was statistically significant (P<10^−4^; t-test; **Figure 5F**). By contrast, pairs of regions that are unconnected in the structural covariance network (at 10% edge density) showed no significant difference between levels of HSE gene co-expression and whole genome co-expression.

### Effects of spatial distance on structural covariance and gene co-expression

A linear function of the logarithmic inter-regional distance value explained 16% of the variance in strength of structural covariance (**Figure S5**). HSE gene co-expression explained 9% of the variance in strength of structural covariance; and cytoarchitectonic class membership explained 6% of the variance. Linear multiple regression demonstrated that the combination of the logarithmic distance, HSE gene co-expression and cytoarchitectonic classification collectively explained 22% of the variance in strength of structural covariance.

## DISCUSSION

Here we have compared the organization of a structural covariance network, derived from cortical thickness measurements in 152 left-hemisphere regions in 296 healthy young participants, to the organization of a transcriptional network, derived from gene expression measurements in the same 152 regions in 6 adult post mortem brains. Since transcriptionally and histologically similar brain regions are more likely to be anatomically connected (Goulas et al. 2016), and since structural covariance is a putative marker of anatomical connectivity, we made two, related hypothetical predictions: i) that co-expression of genes should be correlated with the strength of structural covariance between regions; and ii) that the topology of the structural covariance network should be coupled to the topology of the transcriptional brain network. Considering co-expression of the nearly complete genome, we found no more than modest support for these predictions: the two networks had qualitatively similar topology, their edge weights were significantly correlated and there was evidence for greater whole genome co-expression between brain regions that were assigned to the same topological module of the SCN’s community structure. There was stronger evidence in support of links between structural covariance or SCN topology and regional co-expression of a much smaller set of (19 HSE) genes, known to be enriched specifically in human supra-granular cortex.

### Gene expression and co-expression in the brain

The neocortex displays a remarkable conservation of gene expression across individuals (Hawrylycz et al. 2015), cortical areas (Hawrylycz et al. 2012) and species (Oldham et al. 2006; Zeng et al. 2012). In accordance with the cortical homogeneity of the transcriptomic profile, we found that most of the inter-regional gene co-expression values were extremely significant. Nevertheless, regional differences in cortical gene transcription are subtle but important. Studies based on non-human animals reveal that different cell types display a robust molecular signature where neurons, oligodendrocytes, astrocytes and microglia tend to up-regulate and down-regulate the expression of specific subsets of genes (Lein et al. 2007; Ko et al. 2013; Grange et al. 2014). This transcriptional signature of cell type-specific genes can be used to define both major compartments and smaller anatomical structures of the mouse brain (Ko et al. 2013; Grange et al. 2014). In humans, the grouping of genes according to their co-expression pattern also reveals a modular structure that distinguishes the major cell classes of the underlying brain tissue (Oldham et al. 2009; Eising et al. 2016). Prior studies of the Allen Human Brain Atlas showed, in agreement with mouse studies, that the gene expression pattern reflects the specialization of primary sensorimotor areas and subdivisions of the frontal lobe (Hawrylycz et al. 2012). In keeping with these results, we found that gene co-expression was significantly related to spatial proximity: two cortical regions that are close together will have more similar gene expression profiles (Lein et al. 2007; Bernard et al. 2012; Hawrylycz et al. 2012, 2015). It has been proposed that the effect of anatomical closeness on transcriptomic similarity reflects phylogenetic and ontogenetic distance between regions (Zapala et al. 2005).

### Transcriptomic brain network construction

Gene co-expression networks reported in previous studies refer to networks where the nodes represent genes and edges denote genetic association across samples in a dataset (for a review, see Fontenot and Konopka, 2014). In this type of network, edges link pair of genes that are over/under expressed in the same tissue. The transcriptomic brain network proposed here describes, on the contrary, transcriptional relationships (edges) between spatially delimited cortical regions (nodes). This makes it possible to compare transcriptomic relationships between brain regions to the human brain connectome based on anatomical measures derived from MRI data.

The method for matching AIBS samples to MRI regions of interest in the present study does not rely on manual interventions and relies on more conservative assumptions about homogeneity of gene expression across short distances in the brain than previous studies. In particular, prior work assigned each MRI region of interest to the anatomical structure (as defined in the AIBS dataset) containing the nearest AIBS sample across all donors and then averaged gene expression from all samples falling within that AIBS anatomical region (Vértes et al. 2016; Whitaker et al. 2016). In contrast, French and Paus (2015) took a more straightforward approach where a sample was mapped into a FreeSurfer region if its MNI152 sample coordinate was inside a FreeSurfer cortical region. However, this required visual inspection due to spatial distortions being introduced by coregistration of donors’ brains to standard MNI152-space. In this work we provide a fully automated method for matching MRI parcellation with Allen Brain Atlas samples based on sample mapping in each donor’s native space in order to take into account inter-individual difference of cortical morphology.

### Whole genome co-expression and structural covariance

Using graph theoretical methods to analyse the topology of the transcriptomic brain network we found evidence for topological segregation (high clustering coefficient and modules) and topological integration (short path length, high global efficiency and a rich club). This was a qualitatively similar profile to the complex topology of the structural covariance network. However, the extent of overlap between whole genome co-expression and structural covariance was limited. For example, there was a significant but weak correlation between edge-wise whole genome co-expression and structural covariance; the strength of structural covariance decayed much faster as a function of physical distance than the strength of whole genome co-expression; and only in the SCN (not TBN) was nodal degree significantly related to mean connection distance. The coupling between whole genome co-expression and structural covariance strengthened somewhat when the analysis was restricted to the 10% of network edges representing the strongest correlations in cortical thickness between regions (**Figure 5**). Analysis of the modular community structure of both networks provided stronger (but still modest) evidence of overlap between the structural covariance network and the whole genome transcriptome. Whole genome co-expression was significantly stronger between pairs of nodes that belonged to the same module of the experimentally estimated SCN compared to pairs of nodes belonging to a null distribution of pseudo-modules, generated by permuting empirical modules on the cortical surface while preserving module size, contiguity and distance.

These results should be considered in the context of some relevant prior studies. (Chen et al. 2011, 2012, 2013) evaluated genetic influences on cortical areal expansion by correlating the individual genotype with the rate of deformation needed to map each subjects’s surface into atlas space. The association between genotype and area expansion was used to define the boundaries of a genetic subdivision of the cortex and depict its genetic patterning. Resulting genetic organization revealed a hierarchical, modular configuration consistent with specialized functional and lobar subdivisions. Moreover, this genetic subdivision followed closely the developmental changes of cortical thickness, with age-related cortical shrinkage trajectories varying as a function of inter-regional genetic similarity (Fjell et al. 2015). Cortical thinning in adolescent has been recently associated with the expression of genes coding for glucocorticoid and androgen receptors (Pui-Yee Wong et al. 2017). Genetic associations among regional measures of cortical thickness have been previously described by Docherty et al. (2015). This twin-based study revealed that genetic relationships between structurally-determined regions exhibit small-world properties (Docherty et al. 2015). Fulcher and Fornito (2016) showed that transcriptomic correlations are stronger between hub nodes than between non-hubs of the mouse brain connectome.

### HSE gene co-expression and structural covariance

For the HSE gene set, we did not construct a transcriptional network, and we did not perform topological analyses but rather concentrated on the relationships between HSE gene co-expression, structural covariance and SCN topology. HSE gene co-expression was significantly correlated with structural covariance; high degree hubs in the structural covariance network had higher levels of HSE gene co-expression than less central, low degree nodes; and nodes that belonged to the same SCN module or the same cytoarchitectonic class had higher levels of HSE gene co-expression than pairs of regions that belonged to different modules or classes. These results constitute stronger evidence that structural covariance and network topology is linked to gene co-expression and are arguably compatible with what is known about HSE genes. HSE genes are overexpressed in the supragranular layers (II and III) of the cerebral cortex in humans, but under-expressed in the same layers of mouse cortex (Zeng et al. 2012). This small but substantial species-differential expression is hypothesized to drive the shift from predominantly cortico-subcortical connectivity in non-primate mammals to the major emphasis on cortico-cortical projections in the human brain (Zeng et al. 2012). Convergently, two prior fMRI studies in humans have shown that HSE gene (co)-expression is related to functional connectivity between cortical areas. Krienen et al. (2016) reported stronger HSE transcriptional similarity within than between resting state fMRI networks, which share many similarities to modules; likewise, Vértes et al. (2016) showed that HSE genes are significantly over-represented among the genes that are most over-expressed in cortical areas with high inter-modular degree and long mean connection distance. The data reported here add to this literature by demonstrating for the first time that HSE gene co-expression is also linked to MRI measures of structural covariance and brain anatomical network topology. Thus the evidence is growing that HSE genes play an important role in the formation of human connectome topology, especially high degree hubs and long distance connections. It would be interesting in future studies to explore the effects of sequence variation in HSE genes on human connectome phenotypes and their development.

### Methodological issues

First, it is important to note that the gene expression dataset provided by AIBS included six left hemispheres but only two right hemispheres, due to the expression similarities between hemispheres (Hawrylycz et al. 2012). Given the subsampling of the right hemisphere, we pooled all the samples into the left hemisphere and, consequently, only MRI data from the left hemisphere was included in the analyses. Thus our results cannot be generalized to whole brain or inter-hemispheric connectivity. Third, the tissue samples used for mRNA sequencing were not homogenously distributed across the cortex. As a result, gene expression of each cortical region was calculated by averaging a different number of AIBS samples, leading to a variable signal-to-noise ratio across regions. Fourth, AIBS data were based on 5 male donors and one female, with a mean age of 42.5 years, whereas the MRI data were collected from 296 healthy gender-balanced subjects with a mean age of 19.1 years. Age- and gender-related changes in brain gene expression (Berchtold et al. 2008) and structural covariance of cortical thickness (Vasa et al. 2017) as well as inter individual differences may be an important confounding factor when comparing transcriptomic and neuroimaging data.

## Conclusions

Structural covariance based on MRI measurements of cortical thickness, and transcriptomic brain network had similar complex topological properties, showing organizational patters partially, but not completely overlapped. The high degree hubs of structural covariance networks, were coupled specifically to regional expression and co-expression of a set of genes known to be important for long-range cortico-cortical connectivity of the human brain (HSE genes).

## ACKNOWLEDGEMENTS

We thank Gita Prabu, Roger Tait, Cinly Ooi, John Suckling and Becky Inkster for data collection and storage as well as the Allen Institute for Brain Sciences for access to human brain gene expression data (© 2010 Allen Institute for Brain Science. Allen Human Brain Atlas. Available from: http://human.brain-map.org/.

